# Exploration-exploitation trade-off is regulated by metabolic state and taste value in *Drosophila*

**DOI:** 10.1101/2024.05.13.594045

**Authors:** Samuel C. Whitehead, Saumya Y. Sahai, Jamie Stonemetz, Nilay Yapici

## Abstract

Similar to other animals, the fly, *Drosophila melanogaster,* changes its foraging strategy from exploration to exploitation upon encountering a nutrient-rich food source. However, the impact of metabolic state or taste/nutrient value on exploration vs. exploitation decisions in flies is poorly understood. Here, we developed a one-source foraging assay that uses automated video tracking coupled with high-resolution measurements of food ingestion to investigate the behavioral variables flies use when foraging for food with different taste/caloric values and when in different metabolic states. We found that flies alter their foraging and ingestive behaviors based on their hunger state and the concentration of the sucrose solution. Interestingly, sugar-blind flies did not transition from exploration to exploitation upon finding a high-concentration sucrose solution, suggesting that taste sensory input, as opposed to post-ingestive nutrient feedback, plays a crucial role in determining the foraging decisions of flies. Using a Generalized Linear Model (GLM), we showed that hunger state and sugar volume ingested, but not the nutrient or taste value of the food, influence flies’ radial distance to the food source, a strong indicator of exploitation. Our behavioral paradigm and theoretical framework offer a promising avenue for investigating the neural mechanisms underlying state and value-based foraging decisions in flies, setting the stage for systematically identifying the neuronal circuits that drive these behaviors.

## INTRODUCTION

In their natural habitat, animals must make several behavioral decisions to optimize their fitness and survival. One such behavioral decision that is immensely crucial for the survival of any species is foraging.^1,2^ Foraging is a complex behavior where animals search for food by combining information from their external senses with their body’s metabolic needs.^3–6^ Once a food source is found, the animal must evaluate the current food option based on past experiences and current metabolic state and decide whether to commit to the currently available choice or to explore alternatives at the expense of energy and time.^7–12^ For example, grasshoppers develop a preference for odors they experienced when they were food-deprived compared to odors they experienced during satiety.^13^ Cuttlefish prefer to prey on large live shrimp when hungry, in contrast to choosing smaller prey when they are satiated.^14^ When deprived of sodium, rats consistently seek sodium salts over non-sodium salts (potassium, calcium, etc.)^15^, reducing their preference for other nutrients.^16^ How does the nervous system regulate such foraging decisions? What are the behavioral computations and neural mechanisms animals use to explore their environment and evaluate the available food sources? To study foraging decisions, it is critical to utilize experimental and theoretical approaches together to formulate a framework that explains the foraging strategy by incorporating the metabolic state of the nervous system and the current choices in the animal’s environment.^2,12,17–19^

The fly, *Drosophila melanogaster*, is an emerging model organism for deciphering neural circuits that regulate foraging behavior.^4,10,20,21^ Flies use taste neurons in the chemosensory bristles of the legs, the labellum, and the pharynx to detect and evaluate food sources.^22,23^ Taste neurons express one or multiple members of the chemosensory gene families consisting of the gustatory receptors (Gr)^24,25^, the ionotropic receptors (Ir)^26,27^, the ENaC/DEG, pickpocket (Ppk) channels^28^, and the transient receptor potential (Trp) channels.^29^ Stimulation of sugar-sensitive taste neurons in the labellum or legs triggers extension of the proboscis in fasted flies followed by initiation of food intake.^30–33^ During ingestion, food is sensed by the pharyngeal taste neurons that sustain or terminate the ingestion.^34–36^ Similar to many other animals^37–40^ and humans^41^, flies change their foraging strategy from global search to area-restricted or local search upon encountering a food patch.^21,42–45^ Activation of sugar-sensing taste neurons is sufficient to trigger the local search behavior even in the absence of a food source.^20,46,47^ A recent study suggests that flies use path integration to recall the site of a previously found food source, enabling them to pinpoint its exact location even after some time has passed.^48^ These findings suggest that flies might use a process similar to spatial working memory to modify their foraging strategy and switch between exploration and exploitation behaviors. The neural basis of this mechanism and the impact of metabolic state or taste/nutrient value on the foraging decisions remains poorly understood in flies and other animals.

Here, we developed an automated one-source foraging assay that uses video tracking coupled with high-resolution measurements of food intake to determine how metabolic state and taste/nutrient value impact the foraging behavior of flies. Using our setup, we investigated the behavioral variables flies use when foraging for food with different nutrient values and in varying metabolic states. We found that food or water deprivation can individually and additively stimulate locomotor activity in flies, indicative of elevated foraging behavior. Interestingly, flies lacking all sugar receptors failed to demonstrate a robust local search behavior after locating a food source, suggesting taste sensory information rather than post-ingestive feedback is critical for switching the foraging strategy from exploration to exploitation. Finally, we recorded fly foraging behavior in different hunger states and with varying sugar concentrations to determine which behavioral or state variables flies use to determine their foraging strategy. We found that the dominant factor influencing flies’ proximity to the food source, a strong indicator of exploitation, is their metabolic state and the recent volume of food they ingested. Our behavioral paradigm and theoretical approaches represent a promising tool for investigating the neural basis of state- and value-based foraging decisions in *Drosophila melanogaster.* They can be expanded to other species of insects and maybe even to vertebrates to examine whether universal behavioral strategies are used across species to optimize foraging and food intake decisions.

## RESULTS

### Visual Expresso captures the foraging and feeding decisions of individual flies

Historically, manual behavioral quantifications were used to measure food intake in flies.^49,50^ Recently, multiple automated assays have become available to monitor food intake and foraging behaviors.^51–53^ Previously, we developed the Expresso system to monitor the food ingestion of individual flies in real-time and with high temporal and volumetric resolution.^35^ In this study, we integrated a video tracking system into the Expresso to monitor the foraging strategies of flies as they search for food in a rectangular arena. We named this new system Visual Expresso (V-Expresso). The V-Expresso system is equipped with a machine-vision camera and two Expresso banks consisting of 10 distinct food ingestion recording channels, enabling simultaneous monitoring of the feeding and foraging behavior of 10 individual flies (Figures 1A and 1B, Figures S1A-1C). We also developed a Python-based custom analysis software, Expresso Analysis Toolbox (EAT) (https://github.com/samcwhitehead/EAT), to analyze the Expresso data. EAT has two separate analysis pipelines: The first pipeline analyzes the liquid level data to detect individual feeding bouts of flies automatically. The second pipeline analyzes the camera input to detect individual flies and then track their locomotor activity (Figure 1C). These two analysis pipelines are integrated to validate automated feeding bout detection and to calculate several variables related to fly foraging and feeding behaviors. The behavioral variables we can extract from the V-Expresso dataset include individual feeding bout volume, duration of feeding, total food ingestion volume, time series of feeding bouts, time spent moving in the arena, and the trajectories of flies before and after each food ingestion event. EAT facilitates the analysis of V-Expresso data by providing a user-friendly graphical interface to visualize the individual fly foraging and feeding behaviors (Figures 1D and E). It also supports batch processing of locomotor and ingestion data across groups of flies (Figures 1F-1H). The analysis pipeline for fly feeding bout detection, locomotor tracking, and technical details of the EAT software are explained in the method section and the GitHub webpage. In summary, the V-Expresso system offers a comprehensive platform to study fly foraging and feeding decisions, enabling us to observe changes in their navigation strategies before and after consuming a specific food source in various metabolic states.

**Figure 1.**
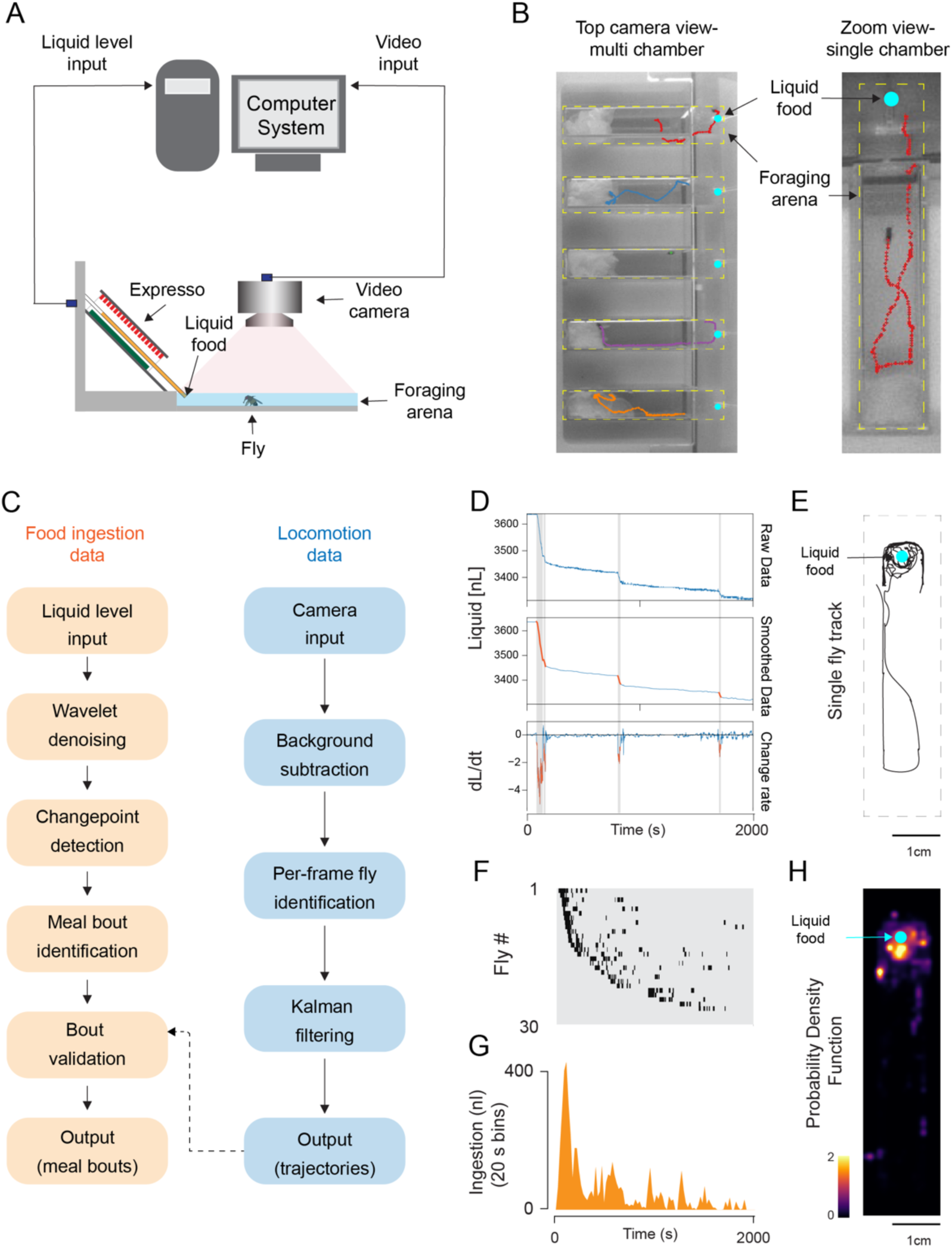
A new behavior system for analyzing foraging behavior in Drosophila. **(A)** Side schematic of the Visual Expresso system. The food intake and locomotion measurements are synchronized using an Arduino board. **(B)** Top view of the Expresso behavior arena (dashed yellow), containing individual flies. Cyan dots indicate the capillary tip where the liquid food is dispensed (left). The dashed red line illustrates an example of a fly’s trajectory (right). **(C)** EAT uses two distinct analysis pipelines for feeding bout and fly trajectory detection. These pipelines are combined to validate feeding bout detection. **(D)** Sequential processing of raw liquid input data and detection of feeding bouts (top to bottom). **(E)** A fly’s trajectory during the entire behavior session. The cyan dot indicates the location of the liquid food. (**F**) A feeding bout raster plot of 30 flies, each row representing data from a single fly, and feeding bouts are indicated by vertical ticks. (**G**) Histogram displaying the total ingestion of all flies tested in 20-second binned intervals. (**H**) Heatmap showing the probability density function of a fly during the behavior session. Example data illustrate the food intake and locomotor activity of a male fly deprived of food for 32 hours and presented with 1M sucrose solution.

### Food and water deprivation stimulate fly locomotion

Using the V-Expresso system, we first investigated how different metabolic states impact the locomotor activity of flies. Animals show increased arousal, motivation, and locomotion during food or water deprivation.^54–56^ In our first set of experiments, we assessed how food deprivation impacted the locomotor activity of flies. We tracked wild-type female flies in a rectangular arena without food or water sources when they were fed or food-deprived for different periods (Figures 2A and 2B). Food deprivation did not change the flies’ walking speed but increased their fraction of time walking and cumulative distance walked (Figures 2C and 2E). We especially observed a significant change in locomotor activity after 16 hours of food deprivation (Figures 2D and 2E). Next, we investigated whether water or food deprivation impacts locomotor activity differently. In these experiments, flies were either 6 hours water-deprived or 24 hours food-deprived or were exposed to both deprivation regimes simultaneously (Figure 2F). Water deprivation had similar stimulatory effects on locomotor activity; thirsty flies walked at a similar speed as food-deprived flies (Figure 2G). The cumulative distance walked was also not different between thirsty or food-deprived flies (Figure 2G). Interestingly, when flies were exposed to both deprivation regimes simultaneously, they walked faster and longer, leading to an increase in cumulative distance walked compared to flies exposed to only food or water deprivation regimes (Figures 2G and 2H). These findings suggest that water or food deprivation can independently enhance the locomotor activity of flies, with their combined effects being additive.

**Figure 2.**
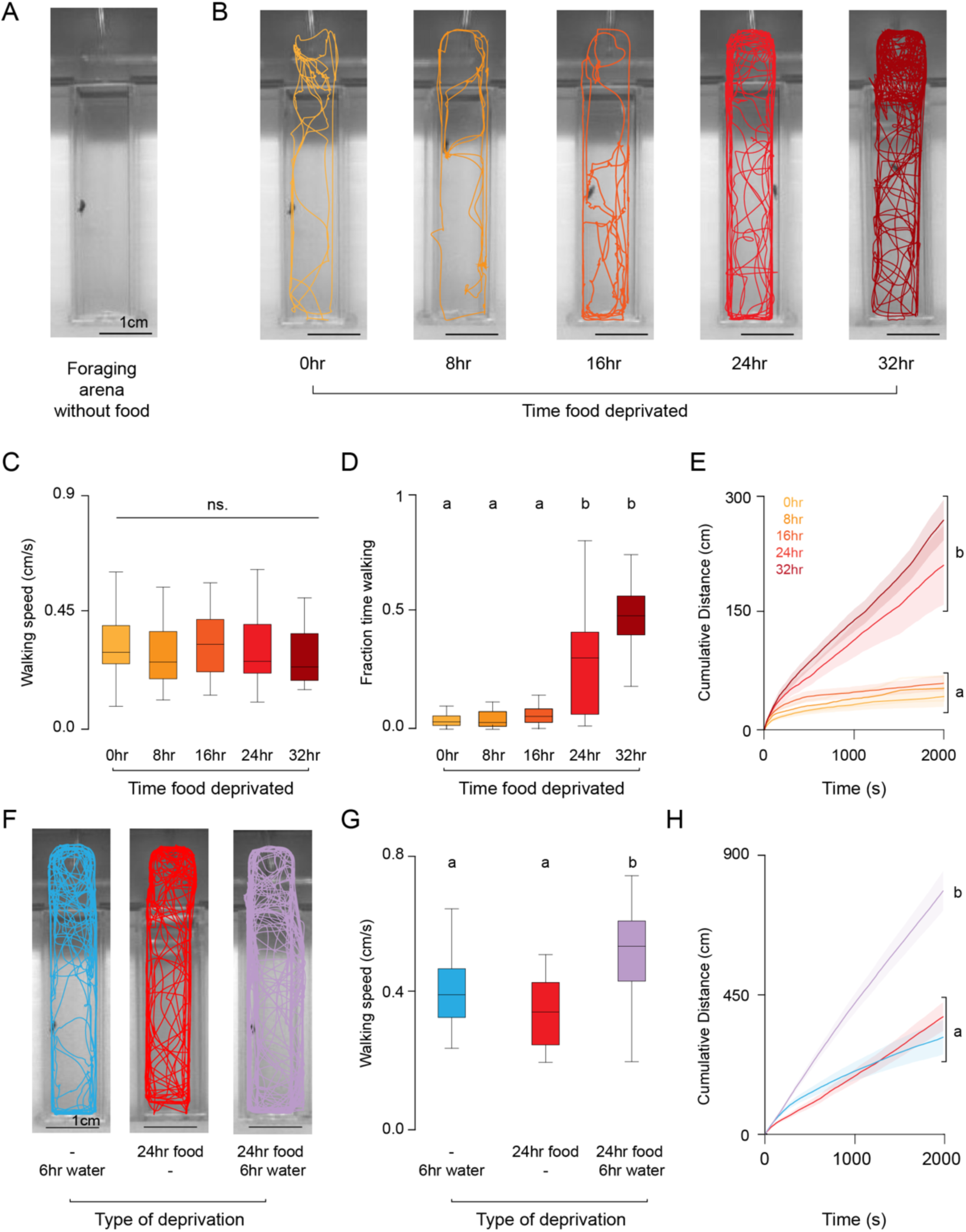
Food and water deprivation stimulate locomotor behavior in flies. **(A)** Top view image of a Visual Expresso behavior arena. (**B**) Walking trajectories of representative wildtype (CS) flies fed ad libitum (0hr) or food deprived for specific durations ranging from 8 to 32 hours, superimposed onto the behavioral arena. **(C)** Average walking speed of flies during locomotion in specified metabolic states (Kruskal-Wallis test, p=0.39, n=29-30). **(D)** Fraction of time spent walking by flies in specified metabolic states (Kruskal-Wallis test with Dunn’s multiple comparisons, p<0.05, n=29-30). **(E)** Cumulative distance walked by flies in specified metabolic states during the total course of the behavior session (Time series permutation test, p<0.05, n=29-30). (**F**) Walking trajectories of representative flies that are water (6hr) or food deprived (24hr) or exposed to both treatments superimposed onto the behavioral arena. **(G)** Average walking speed of flies during locomotion in specified metabolic states (Kruskal-Wallis test, p<0.05, n=30). **(H)** Cumulative distance walked by flies in specified metabolic states during the total course of the behavior session (Time series permutation test, p<0.05, n=30).

### Food deprivation and taste/nutrient value impact feeding and foraging strategy

Many animals exhibit apparent changes in temporal dynamics of food intake and foraging depending on their metabolic state and the quality of the food available. Rats and mice, when deprived of food, engage in longer bursts of licking with elevated sucrose concentrations.^57,58^ Protein-deprived flies choose food patches enriched in amino acids, and satiated flies explore more globally.^10^ To examine the dynamics of food intake and foraging in flies in response to changes in hunger state and food quality, we used the V-Expresso system. We exposed flies to various fasting periods (0, 8, 16, 24, and 32 hours) and subsequently assessed their consumption of 1M sucrose. We observed that as the duration of fasting increased, more flies began to ingest food. (Figures 3A and 3B). Food deprivation enhanced both the total volume consumed and the number of ingestion bouts per fly, although it did not affect the volume of individual bouts (Figures 3C and 3E).

**Figure 3.**
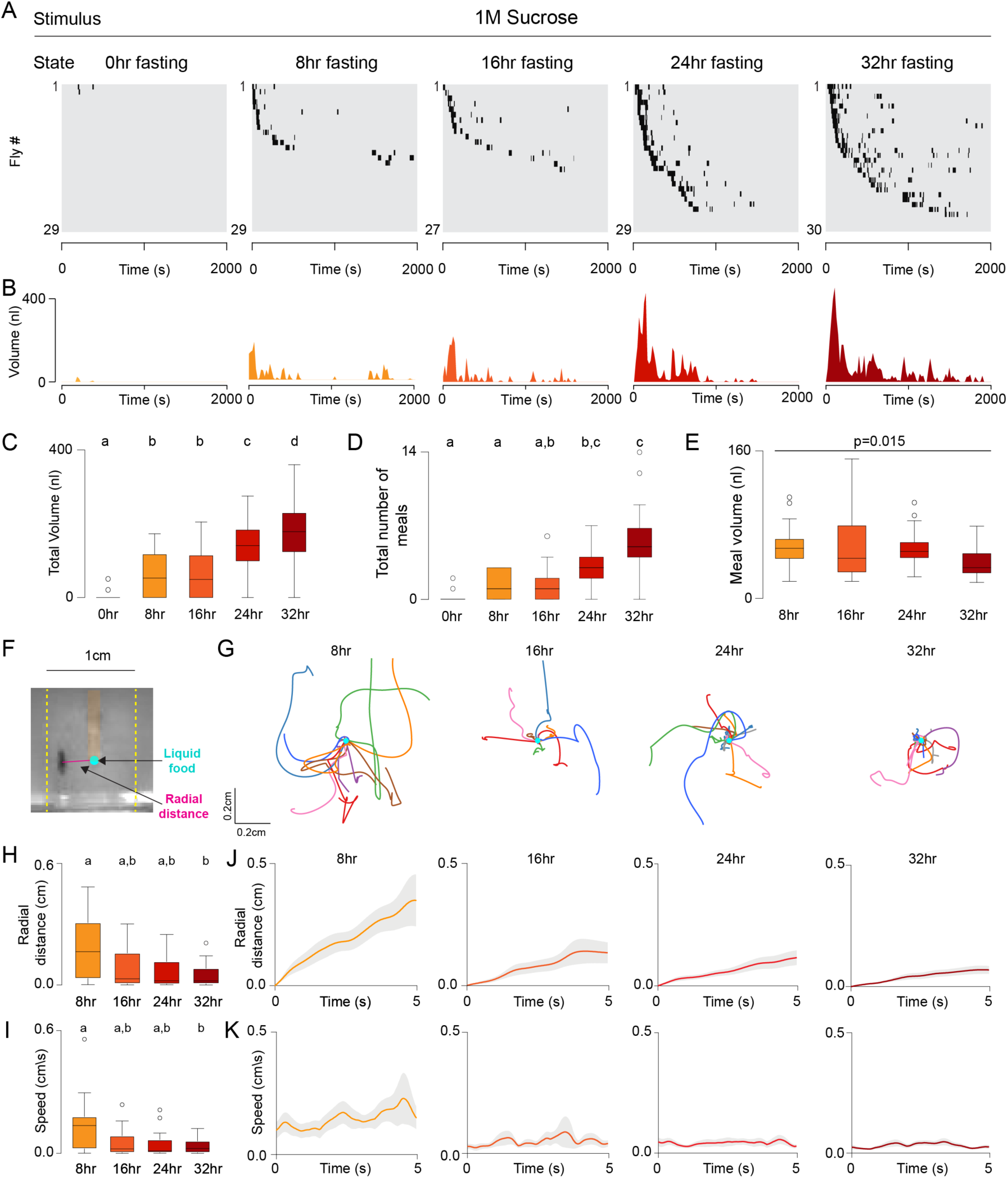
Food deprivation stimulates food intake and exploitation. **(A)** Meal-bout raster plots of wildtype (CS) flies that are fed ad libitum (0hr) or food deprived for specific durations ranging from 8 to 32 hours and presented with 1M sucrose. **(B)** Histogram representation (20-second binned intervals) of the food intake data shown in **A.** (**C)** Total volume and **(D)** total number of meals ingested by flies in specified metabolic states (Kruskal-Wallis test with Dunn’s multiple comparisons, p<0.05, n=27-30). **(E)** Average feeding bout volume ingested by flies in specified metabolic states (Kruskal-Wallis test with Dunn’s multiple comparisons, p=0.015, n=27-30). **(F)** The top-view image of the food zone shows a fly next to the liquid food source; the magenta line indicates the radial distance between the food source and the fly. **(G)** Walking trajectories of individual flies during the 5s following the completion of their first feeding bout in specified metabolic states. (**H)** Average radial distance and **(I)** speed of flies 5s following the completion of their first feeding bout (Kruskal-Wallis test with Dunn’s multiple comparisons, p=0.015, n=16-27). (**J)** Radial distance and **(K)** speed of flies 5s following the completion of their first feeding bout in specified metabolic states.

Next, we investigated the changes in locomotion before and after ingesting 1M sucrose solution. 0-hour fasted flies were not included in this analysis due to the low number of participants in this group that consumed food during the assay time. Upon finding a food source, hungry flies switch their foraging strategy from exploration to exploitation.^21,42,44^ To investigate how the hunger state impacts this switch in foraging strategy, we analyzed the paths of flies at different metabolic states after they finished their first feeding bout. Flies fasted for 8 hours did not switch their foraging strategy; they immediately left the food source after completing their first meal (Figure 3G). However, as fasting time increased, flies decreased their locomotion and walking speed and stayed closer to the food source, indicative of exploitation (Figure 3G). We compared the differences in flies’ foraging strategies at each hunger state by comparing their radial distance to the food source (Figure 3F) and their average speed within five seconds after they finished their first meal. Flies that fasted for 32 hours stayed closer to the food source and moved slower compared to flies that fasted for 8 hours (Figures 3H-3K). Our results suggest that the hunger state is a critical factor in determining flies’ closeness to the food source and switching their foraging strategy from exploration to exploitation.

We next investigated how food quality influences flies’ foraging and feeding strategies. During fasting, animals must maximize their nutrient intake to offset the energy shortfall, which is fundamental to their fitness and survival.^59^ To investigate how food quality impacts fly behavior, we measure the ingestion and foraging dynamics of 32-hour fasted flies that are offered different concentrations of sucrose solutions ranging from 1mM to 1M. At all sucrose concentrations tested, the proportion of flies that consumed food was the same, while the dynamics of food ingestion and foraging strategy differed (Figure 4A). Flies ingested the most significant volume of food at a 100mM sucrose solution and the smallest at 1mM sucrose solution. Interestingly, the total volume of food consumed at 10mM and 1M sucrose was not significantly different (Figure 4C). These results indicate that rapid post-ingestive satiety signals might limit food intake at high sugar concentrations. Sugar concentration also changed the microstructure of ingestion (Figure 4B). Flies consumed fewer bouts from 1mM and 1M sucrose solutions compared to 10mM and 100mM sucrose solutions (Figure 4D). Furthermore, there was a significant change in individual feeding bout volume when flies were offered 1mM or 10mM sucrose compared to 100mM or 1M sucrose solutions; flies consumed larger feeding bouts from higher concentrations of sucrose solutions compared to lower concentrations (Figure 4E). These results indicate that food concentration impacts the temporal dynamics of ingestion, particularly the duration of each feeding bout. Finally, we investigated how sugar concentration impacts foraging behavior. Hungry flies that consumed 1M sucrose decreased their walking speed and stayed closer to the food, indicative of switching from exploration to exploitation behavior. Meanwhile, flies that consumed lower concentrations of sucrose (1mM to 100mM) dispersed away from the source after completing their first meal (Figures 4F-4J). Our findings indicate that flies can optimize their feeding and foraging behaviors according to their hunger level and the quality of available food in their environment.

**Figure 4.**
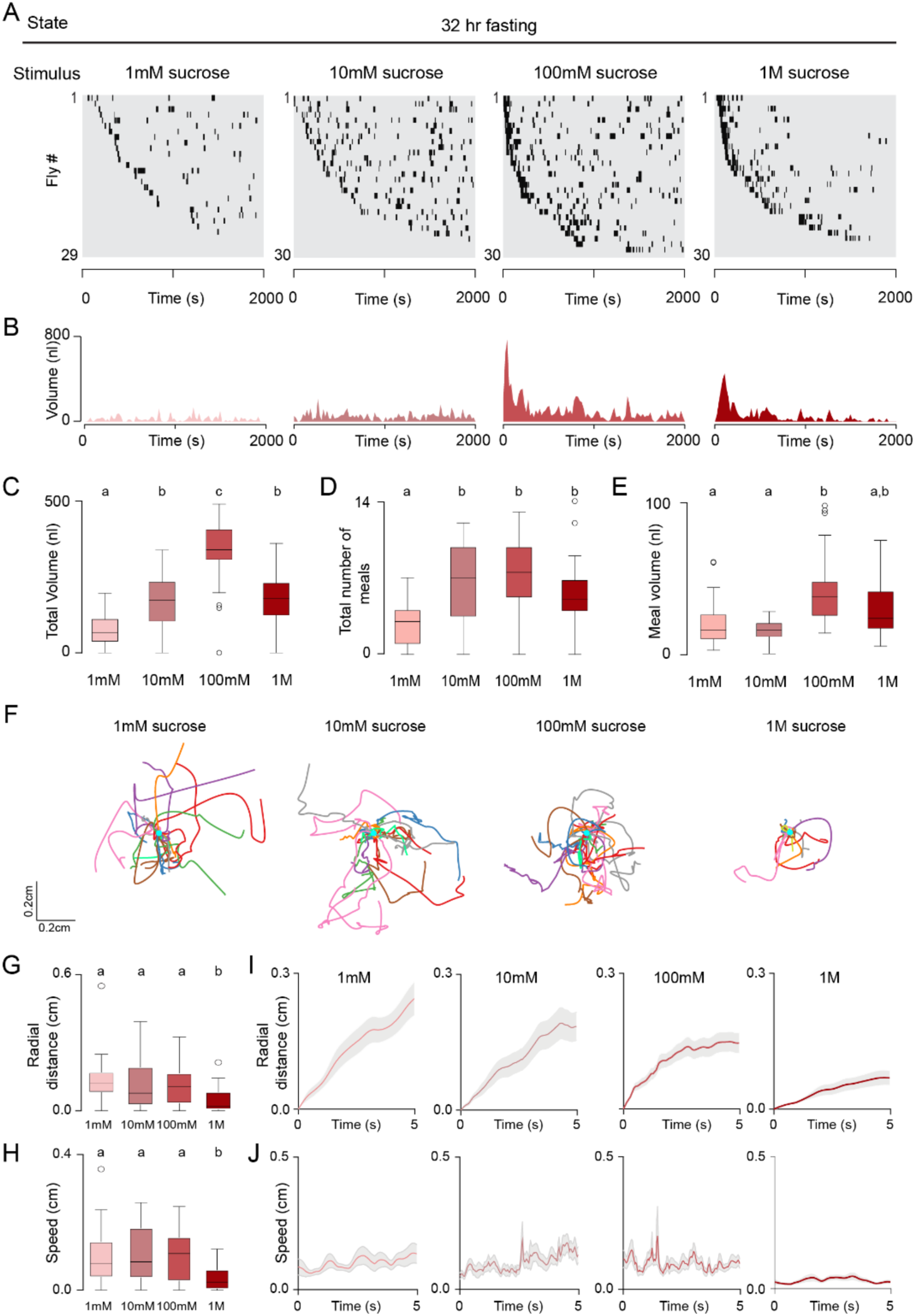
Nutrient value alters exploration-exploitation trade-off in food-deprived flies. **(A)** Meal-bout raster plots of 32-hour food-deprived wildtype (CS) flies presented with sucrose solution ranging from 1mM to 1M. **(B)** Histogram representation (20-second binned intervals) of the food intake data shown in **A.** (**C)** Total volume and **(D)** total number of meals ingested by flies presented with sucrose solution ranging from 1mM to 1M (Kruskal-Wallis test with Dunn’s multiple comparisons, p<0.05, n=29-30). **(E)** Average feeding bout volume ingested by flies presented with sucrose solution ranging from 1mM to 1M (Kruskal-Wallis test with Dunn’s multiple comparisons, p<0.05, n=29-30). **(F)** Walking trajectories of individual flies during the 5s following the completion of their first feeding bout from the indicated sucrose solutions (**G)** Average radial distance and **(H)** speed of flies 5s following the completion of their first feeding bout (Kruskal-Wallis test with Dunn’s multiple comparisons, p<0.05, n=25-29). (**I)** Radial distance and **(J)** speed of flies 5s following the completion of their first feeding bout from the given sucrose solutions.

### Sugar receptors mediate ingestion and foraging behavior

Flies detect food using gustatory receptors expressed in taste neurons distributed across the labellum, wings, legs, and internal mouthparts.^22,23^ Nine gustatory receptors (Gr5a, Gr43a, Gr61a, and Gr64a-f) are shown to detect various types of sugars.^33^ Sugar-sensitive Gr(s) are phylogenetically related and expressed in overlapping taste sensory neuron subsets, suggesting they might function as a heteromeric taste receptor complex.^23^ Gr43a, the fructose receptor, is expressed not only in peripheral taste organs but also in central brain neurons, where it senses circulating fructose.^60^ Since food concentration impacts fly foraging (Figure 4), we hypothesized that gustatory receptor mutants might have defects in switching from exploration to exploitation upon ingesting food. To test our hypothesis, we measured the sugar ingestion and foraging behaviors of flies lacking sugar receptors in the V-Expresso (Figures 5A and 5B). exhibited ingestive behaviors similar to those of control flies. In contrast, flies lacking all sugar receptors (sugar-blind flies) showed a decrease in both the, total volume of sucrose ingested and the total number of meals consumed (Figures 5C and 5D). The meal volume ingested was also at similar levels across sugar receptor mutants except for the sugar-blind flies (Figure 5E). Interestingly, flies lacking Gr43a (ΔGr43a) showed an increase in meal volume when compared to control flies (Figure 5E); this difference was especially evident in the first bout consumed (Figure 5F). Gr43a is expressed not only in taste neurons but also in nutrient-sensitive neurons in the central brain.^60^ Our results are consistent with prior studies demonstrating Gr43a as a nutrient sensor that restricts sugar intake.^61^

**Figure 5.**
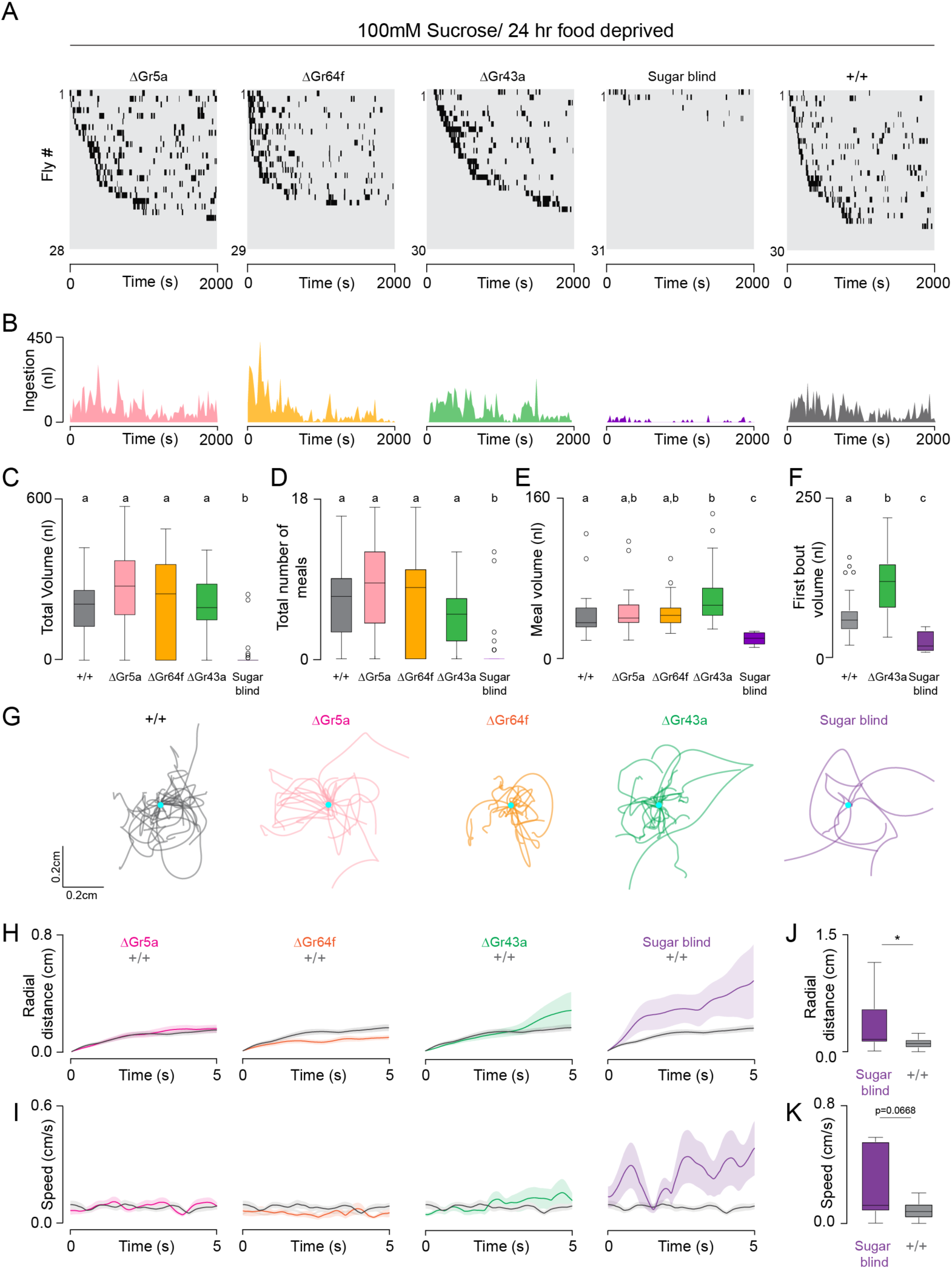
Flies lacking all sugar receptors fail to switch to exploitation. **(A)** Meal-bout raster plots of 24-hour food-deprived flies with indicated genotypes presented with 100mM sucrose solution. **(B)** Histogram representation (20-second binned intervals) of the food intake data shown in **A.** (**C)** Total volume and **(D)** total number of meals ingested by flies with indicated genotypes presented with 100mM sucrose solution (Kruskal-Wallis test with Dunn’s multiple comparisons, p<0.05, n=28-31). **(E)** Average feeding bout volume ingested by flies presented with 100mM sucrose solution (Kruskal-Wallis test with Dunn’s multiple comparisons, p<0.05, n=28-31). **(F)** First feeding bout ingested by flies with indicated genotypes (Kruskal-Wallis test with Dunn’s multiple comparisons, p<0.05, n=28-31). **(G)** Walking trajectories of individual flies with indicated genotypes during the 5s following the completion of their first feeding bout. (**H)** Radial distance and **(I)** speed of flies with indicated genotypes after consuming their first feeding bout from the 100mM sucrose solution. (**J)** Average radial distance and **(K)** speed of flies with indicated genotypes 5s following the completion of their first feeding bout (Mann-Whitney non-parametric, p<, n=7-26).

Next, we investigated how the absence of sugar receptors impacts the switch from exploration to exploitation. Flies lacking Gr5a, Gr64f, or Gr43a showed similar locomotor behaviors compared to control flies (Figure 5G). They decreased their walking speed and stayed closer to the food source after finishing their first meal (Figures 5H and 5I). However, sugar-blind flies failed to switch their foraging behavior from exploration to exploitation after consuming their first meal. These mutant flies did not decrease their walking speed after consuming the sucrose solution and dispersed away from the food area immediately after completing their first meal (Figures 5H and 5I). Our analysis showed a significant increase in the average radial distance of sugar-blind flies compared to controls (Figure 5J). While we observed a trend toward faster walking speeds in sugar-blind flies, it was not statistically significant compared to controls (Figure 5K). Our data suggest that different sugar receptors may have overlapping functions in controlling the foraging behavior of flies. Thus, flies only fail to transition from exploration to exploitation when they miss all sugar receptors. These results indicate that taste sensation rather than post-ingestive nutrient feedback is critical for altering flies’ foraging strategies.

### Radial distance from the food source is determined mainly by metabolic state and ingestion volume

Our results suggest that fly feeding and foraging behaviors are under bi-directional control of metabolic state and taste value (Figure 6A). However, we have not explicitly tested how these two variables interact with each other to determine the volume of ingestion and the switch from exploration to exploitation. To quantify the contribution of metabolic state and taste value as well as other factors such as the ingested meal volume and the nutrient value (calories) in regulating fly feeding and foraging strategies, we conducted an experiment where we varied both the fasting duration (0 to 32 hours) and the food concentration (1mM to 1M) in groups of flies and tested their behavior in the V-Expresso (Figure 6B-6D). As expected, we found a dynamic interaction between the metabolic state and the taste value in determining the number of feeding bouts, the total food intake volume, and the calories ingested (Figure 6B). The total number of feeding bouts and the food intake volume increased with longer fasting regimes across all tested sucrose concentrations. Furthermore, increasing sucrose concentration from 1mM to 100mM elevated the number of feeding bouts and the total volume ingested, indicative of a dose-dependent stimulatory effect on feeding behavior. However, this trend was reversed upon a subsequent increase in sucrose concentration to 1M, which reduced the total number of feeding bouts and the volume consumed (Figure 6B left and middle panels). However, the total calorie consumption was still highest at a 1M sucrose concentration (Figure 6B, right panel), suggesting that the inhibitory effect on food intake might be related to a rapid satiety response induced by ingesting 1M sucrose. These results indicate a threshold beyond which higher sucrose concentrations may inhibit food intake and limit ingestion.

**Figure 6.**
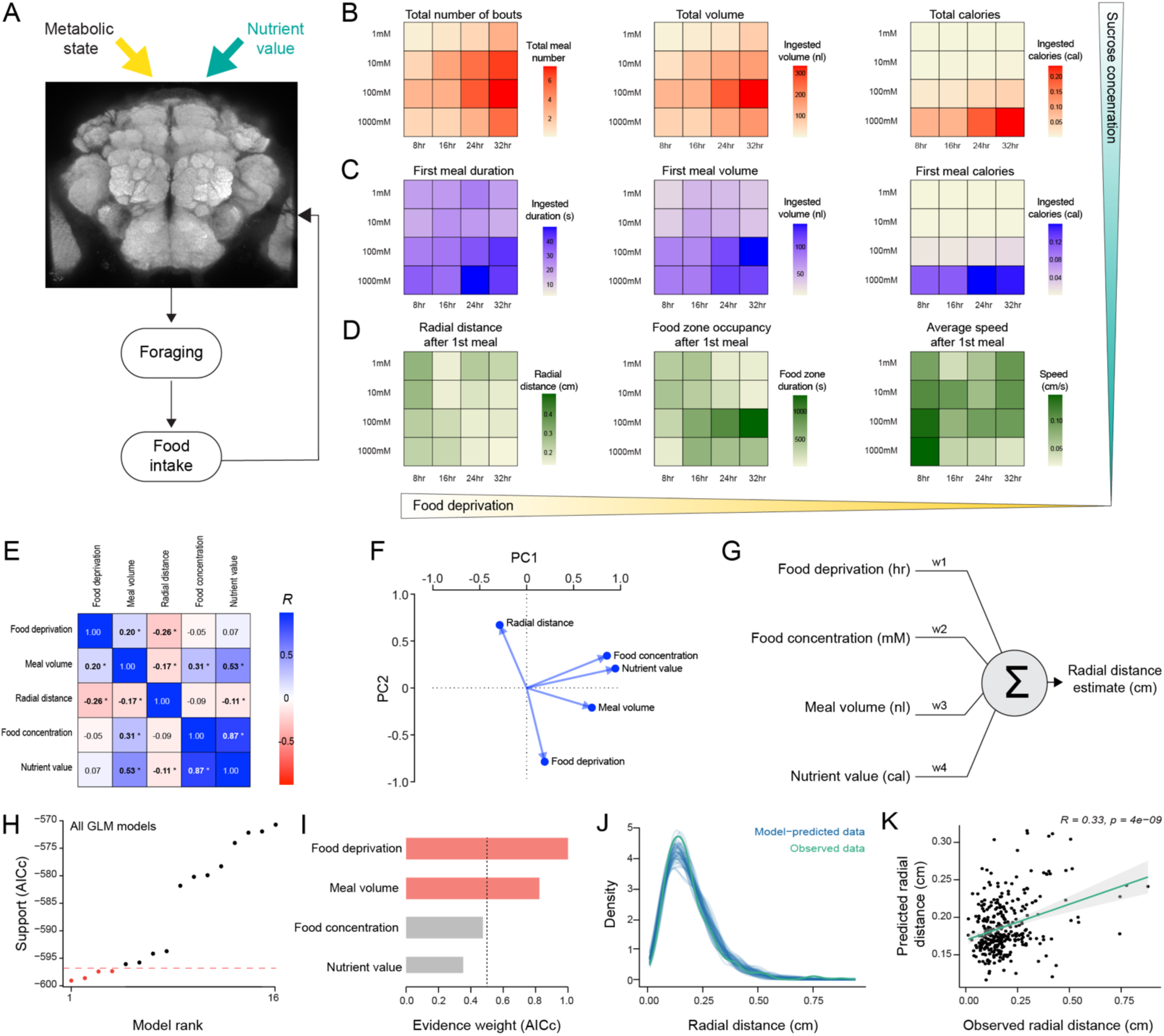
Food deprivation and meal volume are the main variables determining the exploration-exploitation trade-off after food intake. **(A)** Schematic illustrating the integration of metabolic state and nutrient value within the brain to influence foraging and feeding behavior. **(B-D)** Heat maps illustrate the various food intake and foraging variables of flies that are food deprived for durations ranging from 8 to 32 hours and are presented with 1mM to 1M sucrose solutions (n=27-30 per condition, total n=588). **(E)** Heatmap of pairwise correlations between selected food intake and foraging variables. The color key of the correlations is shown on the right. Statistically significant correlations are indicated with a star (Pearson pairwise correlation, R, p<0.05). **(F)** Principal components analysis (PCA) to visualize the relationship between food intake and foraging variables. Two dimensions are shown: PC1 and PC2. **(G)** The schematic of the generalized linear model (GLM). GLM was used to quantify the relationship between food intake and foraging variables (**see Methods**). **(H)** Ranking of GLMs based on AICc, a version of Akaike information criterion (AIC) that has a correction for small sample sizes. The horizontal line represents the AICs limit two units above the best model. **(I)** Estimated weights of model predictors across all GLM models generated. The model predictors selected for the final GLM are colored in red. **(J)** Graphical posterior predictive check comparing the observed distribution of radial distances of flies relative to the food source (green line) to simulated datasets from the posterior predictive distribution (blue lines) **(K)** Scatter plots with linear regression line and the correlation coefficient (Pearson’s R, p<0.05) for observed vs. GLM predicted radial distance values.

Flies alter their foraging behavior after consuming their first meal (Figure 3G and Figure 4F). To investigate how first meal bout impacts fly foraging, we examined its dynamics across different fasting durations and food concentrations. Our data reveal that the duration and volume of the first meal showed similar temporal dynamics to overall food intake (Figure 6B and 6C). Increasing the concentration of the food source, particularly following a period of food deprivation, enhanced the volume of the first feeding bout (Figure S2A), indicating a similar dose-dependent elevation in ingestion as observed in the overall food intake metrics. Additionally, longer fasting regimes increased the duration and volume of the first feeding bout when flies encountered 100mM and 1M sucrose. This increase was not observed when flies were exposed to lower concentrations of sucrose (Figure S2B). These results once again illustrate the dynamic interaction between the metabolic state and the taste value in regulating fly feeding behavior.

After completing our analysis on feeding dynamics, we next investigated how foraging behavior is altered by metabolic state and taste value. We calculated the radial distance of flies five seconds after they had finished their first feeding bout. Five seconds was selected as the time frame based on our observations that within this time frame, flies either decide to stay close to the food source or disperse away from it (Figure 3G and Figure 4F). In these analyses, we found that higher sucrose concentrations and longer fasting durations led to a decrease in the radial distance traveled post-consumption (Figure S3A and S3B) and an increase in the duration of food zone occupancy (Figure 6D middle panel). The food zone occupancy was maximum when flies were fasted for 32 hours and encountered 100mM sucrose solution. We hypothesize that flies had switched their foraging strategy from exploration to exploitation in these conditions. Moreover, there was a noticeable decrease in locomotor activity and walking speed after consuming 1M sucrose solution when flies were food-deprived for longer durations (>8 hours) (Figure 6D right panel). These results are consistent with our previous analysis, which shows that ingestion of 1M sucrose induces a rapid satiety response to inhibit feeding and foraging behavior. Together, our findings illustrate that metabolic state and taste value not only influence immediate feeding decisions but also affect subsequent foraging strategies of flies.

To further characterize the link between feeding and foraging strategies, we next investigated the relationship between metabolic state (food deprivation), taste value (food concentration), and various feeding and foraging variables: meal volume, nutrient value (calories), and radial distance from a food source. We first built a correlation matrix to determine the pair-wise relationship between these variables. We found that food deprivation was positively correlated with meal volume but was negatively correlated with radial distance (Figure 6E and Table S1). On the other hand, food concentration had a positive correlation with both meal volume and nutrient value, but it did not influence radial distance (Figure 6E and Table S1). We further performed principal component analysis on our data set as an independent method to evaluate the relationship between our variables. We found that two principal components, PC1 and PC2, can explain most of the variance in our dataset (Figure 6F). Among the variables tested, food concentration, nutrient value, and meal volume were significant contributors to the variance in the data explained by PC1. In contrast, radial distance and food deprivation impacted PC2 in opposite directions. Together, these results suggest that the feeding and foraging strategies of flies are under the strong influence of the metabolic state; notably, there is a strong negative correlation between flies’ metabolic state and the radial distance to the food source.

To further characterize the contribution of metabolic state and other feeding variables to the flies’ foraging strategy as quantified by its radial distance from the food source, we built a Generalized Linear Model (GLM) (Figure 6G). In the GLM, the radial distance of flies to the food source was selected as the dependent variable, and food deprivation state, food concentration, meal volume, and nutrient value (calories) were selected as predictors. We assumed a gamma distribution (see methods for detailed explanation) and treated the predictors as continuous variables in the model, with no interactions among them. (Figure 6G). We tested several Gamma-GLMs consisting of a combination of our predictors to choose the best-fitted model for our data set (Table S2). The models were ranked using the Akaike Information Criterion corrected (AICc), with the best predictive model being the one with the lowest (most negative) AIC value.^62^ We identified four models with acceptable AICc scores for predicting the radial distance of flies (Figure 6H and Table S2). In these models, food deprivation and meal volume were most influential on the model’s predictive power (Figure 6I). In fact, the model that best predicted the radial distance from the food source used only two variables: metabolic state (food deprivation) and meal volume (Table S2). Next, we tested our model’s efficiency by comparing the observed data to the model’s predictions (Figure 6J). We found a convincing overlap between the density plots of observed and predicted data (Figure 6J). Furthermore, correlation analysis showed a moderate but highly significant positive linear relationship (R=0.33, p=4e-09) between observed and predicted radial distances, reinforcing the model’s validity (Figure 6K). Together, these results indicate that flies’ radial distance, a proxy for their foraging strategy, can be predicted by their metabolic state and ingested food volume.

## DISCUSSION

In this paper, we developed and used V-Expresso, an automated system that combines real-time video tracking and detailed volumetric analysis, to study fly feeding and foraging behaviors. We found that flies’ feeding and foraging strategies are under bi-directional control of their metabolic state and the nutrient environment they encounter. Flies lacking all sugar receptors failed to change their foraging strategy from exploration (searching for food) to exploitation (consuming food) upon discovering a food source, indicating that taste value is a critical factor that drives the foraging strategy of flies. We built a Gamma-GLM model incorporating flies’ metabolic state and ingested meal bout volume to predict their radial distance from the food source. Our experimental and theoretical framework provides a robust platform for identifying molecular pathways and neural circuits that regulate the temporal dynamics of feeding and foraging in flies and for uncovering the dynamic neural computations underlying these behaviors in the fly brain.

### Impact of metabolic state on locomotor activity

In many species of animals, nutrient deprivation induces a state of high locomotor activity.^42,63–65^ Here, we demonstrate a clear impact of metabolic states, hunger and thirst, on the locomotor activity of flies. In our experiments, we found that long food deprivation regimes significantly increased the duration and distance walked in the V-Expresso, but not necessarily walking speed. We especially observed a dramatic change in locomotor activity when flies were food-deprived for 24 and 32 hours (Figures 2D&2E). This suggests a heightened state of arousal and motivation in search of food, supporting previous studies that link metabolic deficits to increased locomotion and hyperactivity in flies.^42,63,66^ Additionally, we demonstrated that water deprivation had similar stimulatory effects on the locomotor activity of flies (Figures 2G&2F), as previously observed in other experimental settings.^67^ Interestingly, when flies were subjected to dual deprivation (food and water), a synergistic effect was observed, resulting in increased walking speeds and distances (Figure 2H). These results indicate that different nutrient deficiencies might generate additive motivational drives within the nervous system to enhance the exploratory drive of flies and increase their chances of finding food or water in their environments. Our results align with previous studies on how metabolic stress impacts animal behavior.^68–71^

### Influence of food deprivation and taste value on feeding and foraging behavior

In this study, our experimental setup, V-Expresso, allowed us to investigate how food deprivation and taste value impact the temporal dynamics of feeding and foraging behaviors, such as the volume and frequency of ingestion events. We demonstrated that the metabolic state of flies mainly increases the number of feeding events rather than influencing the temporal dynamics of ingestion (Figure 3C-3E). Flies showed more persistent ingestion bouts while drinking higher concentrations of sucrose and when they were exposed to more extended food deprivation regimes (Figure 4E). Moreover, varying sucrose concentrations provided further insights into how taste and nutrient value influence feeding decisions. We observed a consistent increase in the frequency and volume of feeding bouts until 100mM sucrose concentration, but this trend reversed when the concentration was further increased to 1M sucrose (Figures 4C and 4D). These findings suggest that while taste exerts a stimulatory influence on ingestion, there is a threshold after which post-ingestive signals likely constrain feeding behavior through rapid satiation, which diminishes subsequent food intake. This behavior reflects an adaptive strategy to maximize caloric intake during food scarcity while avoiding over-feeding. In these experiments, we also noticed that metabolic state and taste value also impacted the foraging strategy of flies post-ingestion. Flies that consumed 1M sucrose at shorter deprivation times dispersed from the food source upon completing their first feeding bout, and they did not conduct frequent revisits. In contrast, flies exposed to more extended deprivation regimes stayed close to the food source (Figure 3G). These two locomotor states imply the existence of divergent foraging strategies, likely orchestrated by the fly nervous system: exploration vs. exploitation. We hypothesized that when nutrient deficiency reaches a certain threshold, flies increase their feeding behavior while maintaining a high exploratory drive. However, as the deficiency becomes more severe, this exploratory drive diminishes, leading the flies to exploit the food source they have already discovered. In our experiments, we further found that the foraging strategy of flies was not only determined by the metabolic state but also by the taste/nutrient value of the food source (Figure 3G and 4F). Flies that consumed a poor source of food (1mM or 10mM sucrose) did not switch their locomotor state to exploitation, similar to flies that were food-deprived for shorter durations. These results indicate that flies’ foraging strategy is not only under the control of their metabolic state but also determined by the taste/nutrient value of the food source. An interesting aspect of our results is that although the marginal value of the sucrose source is always very high relative to the rest of the environment that the fly has access to, flies still leave the food source if they are not hungry or if the food source is suboptimal. These findings suggest that flies’ decisions are not solely based on the immediate availability of resources. Instead, these decisions appear to be influenced by an intrinsic assessment of resource quality, which relies on an absolute scale rather than determined by the availability of current resources.

To discriminate the effect of taste vs nutrient value on feeding and foraging, we further explored the role of specific sugar-sensing gustatory receptors (Gr5a, Gr43a, Gr64f) in mediating these behaviors. We found that flies’ feeding and foraging strategies were altered only when all sugar receptors were mutated (Figure 5G). These findings indicate a redundancy within the fly gustatory system for sucrose detection and align with the hypothesis that diverse sugar receptor genes form a heteromeric receptor complex for detecting various sugar molecules.^72–74^ Furthermore, our data suggests that sensory detection of sucrose rather than post-ingestive nutrient signals appears to drive the switch in feeding and foraging behaviors. Our results are consistent with the previous reports demonstrating that the activation of sugar-sensing taste neurons, even without a food source, is sufficient to alter the foraging strategy of flies.^20,48^

### Integrating Metabolic and Sensory Inputs to regulate Feeding and Foraging

Our findings demonstrate that flies adjust their feeding and foraging strategies based on their metabolic state and the availability of food sources, which points toward the existence of a neural circuit that consists of complex sensory and metabolic feedback mechanisms regulating these behaviors. To systematically investigate the influence of each behavioral and metabolic variable in regulating feeding and foraging strategies, we quantified their interactions and constructed a GLM. Our analysis revealed that the metabolic state and meal volume are the most dominant variables influencing flies foraging strategy post-ingestion. The predictive power of our GLM, validated against observed data, supports the utility of V-Expresso in generating reliable and quantifiable insights into the foraging behavior of flies. We also compared the performance of other simplified or enhanced GLMs and found them to be inferior to our final model, which uses only metabolic state and meal volume as predictors. (Figure 6I). It is not surprising that metabolic state is one of the most influential factors determining flies foraging strategy. However, we were surprised to find out that meal volume rather than nutrient value has a more significant impact on the fly’s proximity to the food source. These findings indicate that post-ingestive nutrient signals may act on a slow time scale, primarily influencing the frequency and the persistency of feeding events rather than directly affecting the foraging strategy of flies on rapid time scales. Previous studies have shown that meal volume is determined by the mechanosensory stretch signals originating from the enteric nervous system, which impact flies’ appetite and limit their food intake.^75^ It is possible that other signals originating from the fly gut influence the foraging strategy of flies and govern the switch from exploration to exploitation behavior. In fact, recent studies have shown that the gut-brain axis regulates food preference and goal-directed behavior in mice.^76–78^

## Conclusion

In summary, we have developed an automated assay, V-Expresso, to quantify the feeding and foraging behaviors of flies and tested how the metabolic state and nutrient environment impact these behaviors. Our results demonstrate that V-Expresso is a valuable tool in the study of feeding behavior in flies, offering new perspectives on how metabolic and sensory inputs govern flies’ foraging strategies. By providing a detailed quantifiable understanding of these processes, our study contributes significantly to the broader field of foraging behavior. It also opens new avenues of research investigating the molecular and neural basis of feeding and foraging strategies in a genetically tractable model organism in response to environmental and physiological changes.

## ACKNOWLEDGMENTS

We thank Andrew Hein and the members of the Yapici Lab for their comments on the manuscript. We thank Azahara Olivia and Ralitsa Todorova for their help in GLM modeling. We acknowledge Bloomington Drosophila Stock Center (NIH P40OD018537) for reagents. This project is supported by the Cornell University Nancy and Peter Meinig Family Investigator Program, the Pew Biomedical Scholar Award, the Alfred P. Sloan Foundation Award, and the National Institutes of Health grant NIH-R35 (5R35GM133698).

## AUTHOR CONTRIBUTIONS

N.Y. conceived the project and designed all the experiments. SS developed the Visual Expresso setup hardware and collected the data in Figures 1-4 & 6. SS conducted this work prior to joining Amazon. SCW developed the EAT analysis software and helped with the behavioral model in Figure 6. JS collected the data in Figure 5. N.Y. analyzed the data, generated the figures, interpreted the results, and wrote the paper based on feedback from JS, SCW, and SS.

## STAR METHODS

### RESOURCE AVAILABILITY

#### Lead contact

Further information and requests for reagents should be directed to and will be fulfilled by the lead contact, Nilay Yapici.

#### Materials availability

- This study did not generate new unique reagents.

#### Data and code availability

- The data supporting this study’s findings are available from the lead contact upon reasonable request.
- All code is available in the publicly accessible GitHub repository (https://github.com/samcwhitehead/EAT).
- Any additional information required to reanalyze the data reported in this paper is available from the lead contact upon reasonable request.

### EXPERIMENTAL MODEL AND SUBJECT DETAILS

#### Flies

*Drosophila melanogaster* flies were maintained on conventional cornmeal-agar-molasses medium at 25°C and 60-70% relative humidity under a 12:12 light: dark (LD) cycle (lights on at 9 AM) and aged 5-7 days. The following lines have been maintained as laboratory stocks or were obtained from Huber Amrein: Canton-S (CS, Yapici Lab stock), w^1118^(Bloomington stock #5905), *Gr5a-LexA*; *Δ43a*/CyO; *Δ61*, *Δ64a-f*/Tm6b (sugar null), *Gr5a-LexA-KI* (*ΔGr5a* mutant), *Gr43a-Gal4-KI* (*ΔGr43a* mutant), *Gr64f-LexA-KI* (*ΔGr64f* mutant). Fly stocks are detailed in the Key Resources Table. The complete genotypes used in each figure are listed in Table S3.

### METHOD DETAILS

#### Visual Expresso Hardware and Data Acquisition

Each Visual Expresso sensor bank comprises a printed circuit board with five linear optical array sensors (TAOS, TSL1406R) comprising 768 photodiodes. A microcontroller connects the system to a computer through a Universal Serial Bus port. When a fly drinks liquid food from a glass capillary, the decrease in the liquid level is detected by a photodiode and is used to calculate instantaneous food ingestion.^35^ The FLIR Blackfly S camera (BFS-U3-13Y3M-C) is used for video acquisition. The videos are recorded at a rate of 30 frames per second. The Expresso data acquisition software generates the hardware trigger given to the camera. Using a trigger-based system, an Arduino board synchronizes the camera and the food intake data acquisition. The video recordings and food ingestion data are synchronized as the data acquisition software triggers the camera on and off.

We used a custom-made chamber fabricated from white Delrin plastic to test flies in the V-Expresso system. The feeding chamber has five compartments, each designed to hold a polystyrene cuvette (VWR # 58017-880) into which a single fly is placed. A sliding door is positioned between the cuvette holding the fly and the capillary tip, allowing synchronization of each trial for all five fly compartments. During each trial, we used two feeding chambers to simultaneously monitor the feeding and foraging behavior of 10 individual flies.

#### Visual Expresso Assay Procedures

##### Food deprivation

We used 3- to 5-day-old male or female CS flies to test the effects of food deprivation on foraging and food intake behaviors. Flies were kept on an ad libitum diet or food-deprived for indicated periods (8, 16, 24, or 32 hours) in vials containing Kimwipes soaked in distilled (milli-Q) water. Food deprivation was started at a time such that the flies reached the appropriate level of food deprivation during their active cycle (3-7 PM). The foraging and food ingestion behaviors were recorded in the Visual Expresso in the absence (Figure 2) or presence of a food source (Figures 3,4 & 6). We used different concentrations of sucrose solution ranging from 1mM to 1M as a food source. The duration of each recording was 2000 seconds. All experiments were run from 3 to 7 PM.

##### Sugar receptor mutants

3- to 5-day-old male flies (*w^1118^* or sugar receptor mutants) were food-deprived on milli-Q water-soaked Kimwipes™ for 24 hours prior to Visual Expresso recordings. We used a 100mM sucrose solution as a food source. The duration of each recording was 2000 seconds. All experiments were run from 3 to 7 PM (Figure 5).

##### Food and water deprivation

We used 3- to 5-day-old female CS flies to test the effects of food and water deprivation on locomotion. Flies were prepared for the experiments as previously described.^79^ Briefly, female flies were either food-deprived on (milli-Q) water-soaked Kimwipes™ for 24 hours or water-deprived on the desiccant for 6 hours with a paper lined with 2M dry sucrose. We also tested a third group of flies that were first food-deprived for 18 hours and then kept on desiccant for an additional 6 hours before the experiments. The duration of each recording was 2000 seconds. All experiments were run from 3 to 7 PM (Figure 2).

### QUANTIFICATION AND STATISTICAL ANALYSIS

#### Visual Expresso Data Analysis Software

Food intake and tracking data collected using the Visual Express system were analyzed using custom software written in Python. Food intake and tracking data are primarily processed in separate pipelines (Figure 1C), the details of which are described below. The Visual Expresso analysis source code and instructions for installation and usage can be found in the GitHub repository. (https://github.com/samcwhitehead/EAT).

##### Food intake data analysis

Raw liquid level time series from the Expresso sensor banks were first denoised using the wavelet-based VisuShrink procedure with soft thresholding according to the universal threshold^80^. Putative feeding bouts were identified via changepoint detection applied to the time derivative of the denoised liquid level data. Specifically, the pruned exact linear time (PELT) method was used to detect changepoints^81^, and a putative feeding bout was defined as an interval between two changepoints in which the median slope of the liquid level data is negative (i.e., decreasing liquid level). Because evaporation from the fluid capillary caused a decrease in liquid level across an experiment, this set of putative feeding bouts included periods of both evaporation and food consumption. To separate these two types of intervals in the time series, the median slope during each putative feeding bout was used to calculate a modified z-score for each bout, *z\** = 0.6745*(*x_i_* − *X*)/MAD, where *x_i_* is the slope of a putative bout, *X* is the median slope across all putative bouts, and MAD is the median absolute deviation across all putative bouts. Putative bouts with a modified z-score >3 were then defined as true feeding bouts, provided the liquid volume consumed and the duration exceeded the experimentally determined noise threshold for meal detection (6nl and 1s, respectively).

##### Video tracking analysis

Video analysis was performed using the OpenCV toolkit.^82^ In brief, videos that simultaneously monitored multiple flies in individual Expresso banks were cropped to produce videos that included only a single fly and an Expresso bank (Figure 1B). These single-fly videos were then processed to subtract the non-uniform background using a mixture of Gaussians model, and the resulting background-subtracted video frames were binarized to identify pixel clusters corresponding to the fly at successive time points. The centroid of these pixel clusters was used to determine the fly’s center of mass at each time point, with a Kalman filter used to both prevent spurious detection events and predict values for the fly’s position when it was occluded from the camera view. Based on the known dimensions of the Expresso bank chamber, the fly center of mass location was converted from units of pixels to centimeters. The 2D fly trajectories were low-pass filtered using a fourth-order Butterworth with a critical frequency of 60Hz to remove tracking noise.

##### Combined analysis

As an additional check on the meal detection process described above, which used only liquid-level data, we imposed criteria for true feeding bouts based on the results of our video tracking. Specifically, during a putative feeding bout, a fly’s median distance from the capillary tip was required to be <0.5 cm in both the x and y direction, and its median speed was required to be <0.1 cm/s. This additional set of conditions was added to the liquid-level-based criteria above to improve further the detection of food intake events (Figure 1C).

#### Statistics

Sample sizes used in this study were based on previous literature in the field. Experimenters were not blinded in most conditions, as almost all data analyses were automated and done using a standardized computer code. All statistical analyses were performed using Prism Software (GraphPad, Version 10.1.1), R-studio (“Desert Sunflower” Release (cd7011dc, 2023-10-16) for macOS), or EAT-data analysis code written in Python (Python 3.7.0). A non-parametric Kruskal-Wallis test followed by Dunn’s multiple comparisons was used to compare more than two genotypes or conditions. The Mann-Whitney non-parametric test was used to compare two genotypes or two conditions. Times series comparisons were done using the time series permutation test in Python. Data labeled with different letters are statistically different.

#### PCA Analysis

We used Principal Component Analysis (PCA) (GraphPad, Version 10.1.1), a multivariate technique, to determine the interactions between the variables we quantified in the Visual Expresso experiments. The following variables were selected for further analysis: the radial distance (RD) from the food source, food deprivation (S), food concentration (C), meal volume (V), and nutrient value (N).

#### GLM Modelling

We used R-studio (“Desert Sunflower” Release (cd7011dc, 2023-10-16) for macOS) packages “glmulti” and “easystats” for data modeling and evaluation. glmulti R package builds all possible Generalized linear models (GLM) with all possible combinations of predictors.^83^ We used a GLM assuming gamma distributed errors with an inverse link function because none of the dependent variables or predictors have a negative value in our data set.^84^ The Akaike Information Criterion (AIC)^85^ is a tool for estimating the prediction error and the comparative quality of different GLMs based on a specific dataset. When presented with multiple models for the same data, the AIC evaluates the quality of each model in comparison to the others.^86^ We used the bias-corrected Akaike information criterion (AICc)^62^ to rank the models generated by glmulti. The AICc score was used because it is correct for the low number of models, and it is less likely to select an over-parameterized model than AIC. GLMs with the lowest AICc scores were chosen for further evaluation (Figure 6H). We focused on the first four models (Table S2) that were less than 2 units away from the GLM with the lowest AICc score and chose the one with the least number of predictors as the final model. In this model, the radial distance (RD) from the food source is the dependent variable. In contrast, food deprivation (S), food concentration (C), meal volume (V), and nutrient value (N) value are predictors. The final GLM equation thus was:

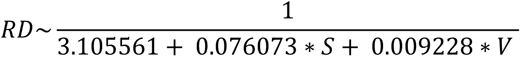

