## Supplementary material for "Exploration-exploitation trade-off is regulated by metabolic state and taste value in *Drosophila*": Table S1

**SUPPLEMENTARY FIGURES and TABLES**

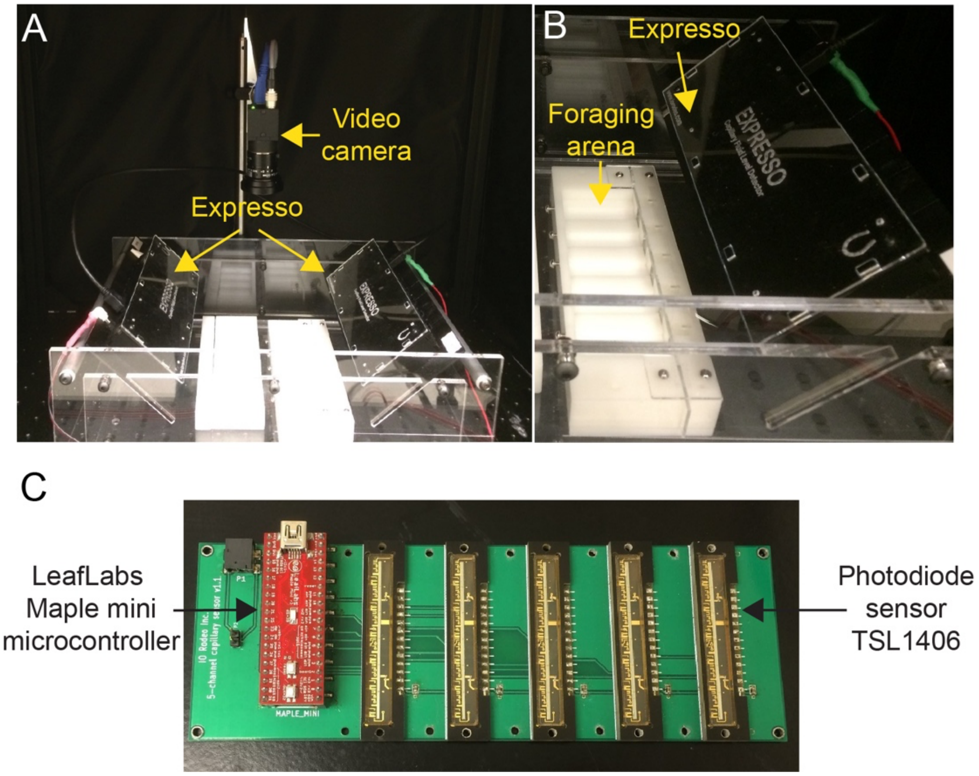


**Figure S1. *V*-Expresso system**

**(A)** Top-down view of the V-Expresso system. A machine vision video camera is used to record the fly's behavior.

**(B)** Side view of the V-Expresso system. The foraging arena is located right below the Expresso food ingestion measurement device. The glass capillaries connect the Expresso to the foraging arena. Flies are allowed to consume food from the glass capillary freely.

**(C)** Inside of the Expresso device. We used a LeafLabs Maple mini microcontroller, to read the data from the TSL1406 photodiode sensors.

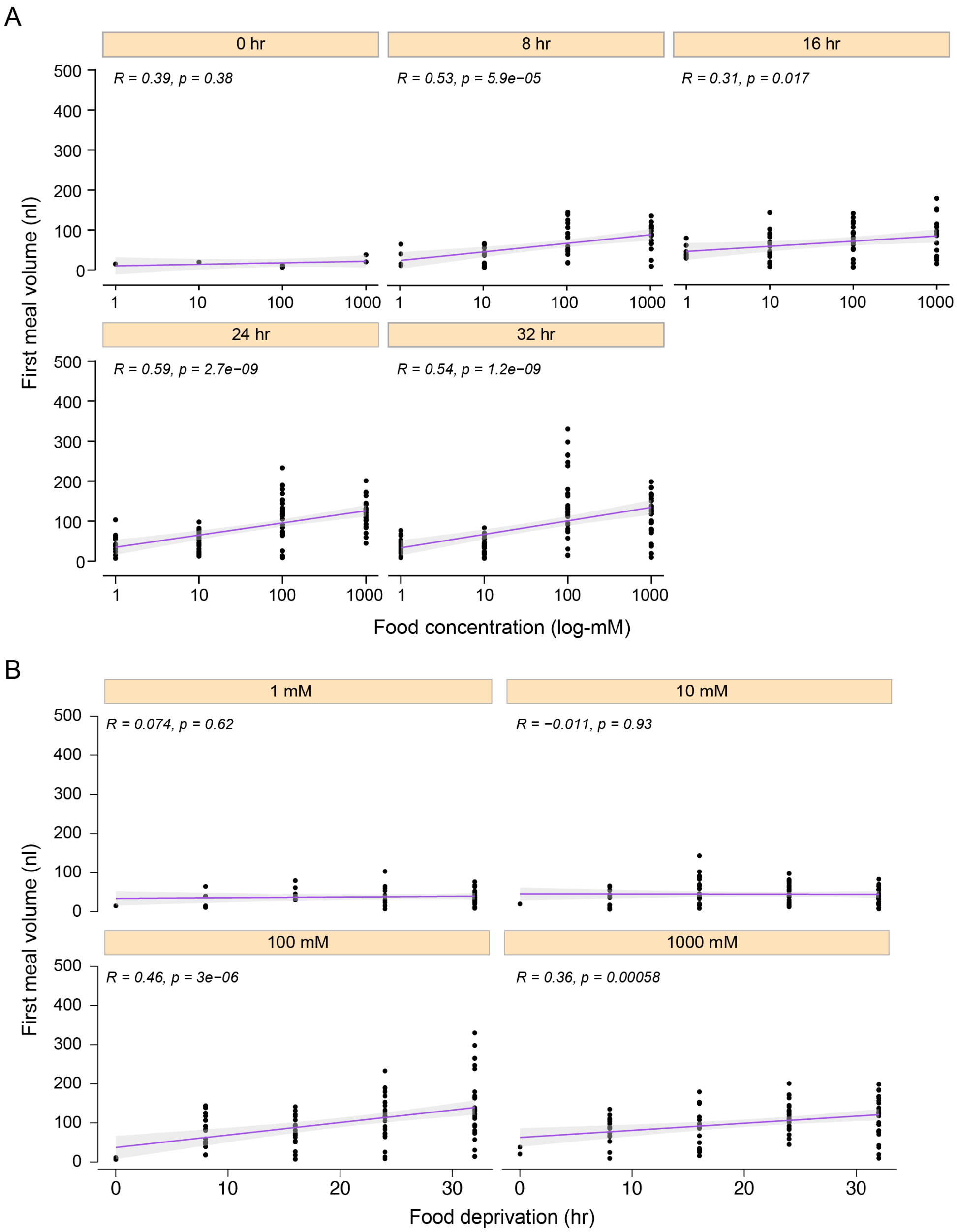


**Figure S2. Metabolic state and taste/nutrient value influence the first ingestion volume**
**(A)** There is a linear correlation between the volume of the first meal and food concentration. Flies fed ad-libitum does not change the volume of their first feeding bout across different sucrose concentrations. In contrast, all flies that have been deprived of food for 8 to 32 hours increase the volume of their initial feeding bout as food concentration rises.
**(B)** There is a linear correlation between the volume of the first meal and food deprivation. This linear relationship is not present at low concentrations of sucrose solution (1mM and 10mM) but is very significant at high concentrations (100mM and 1M).

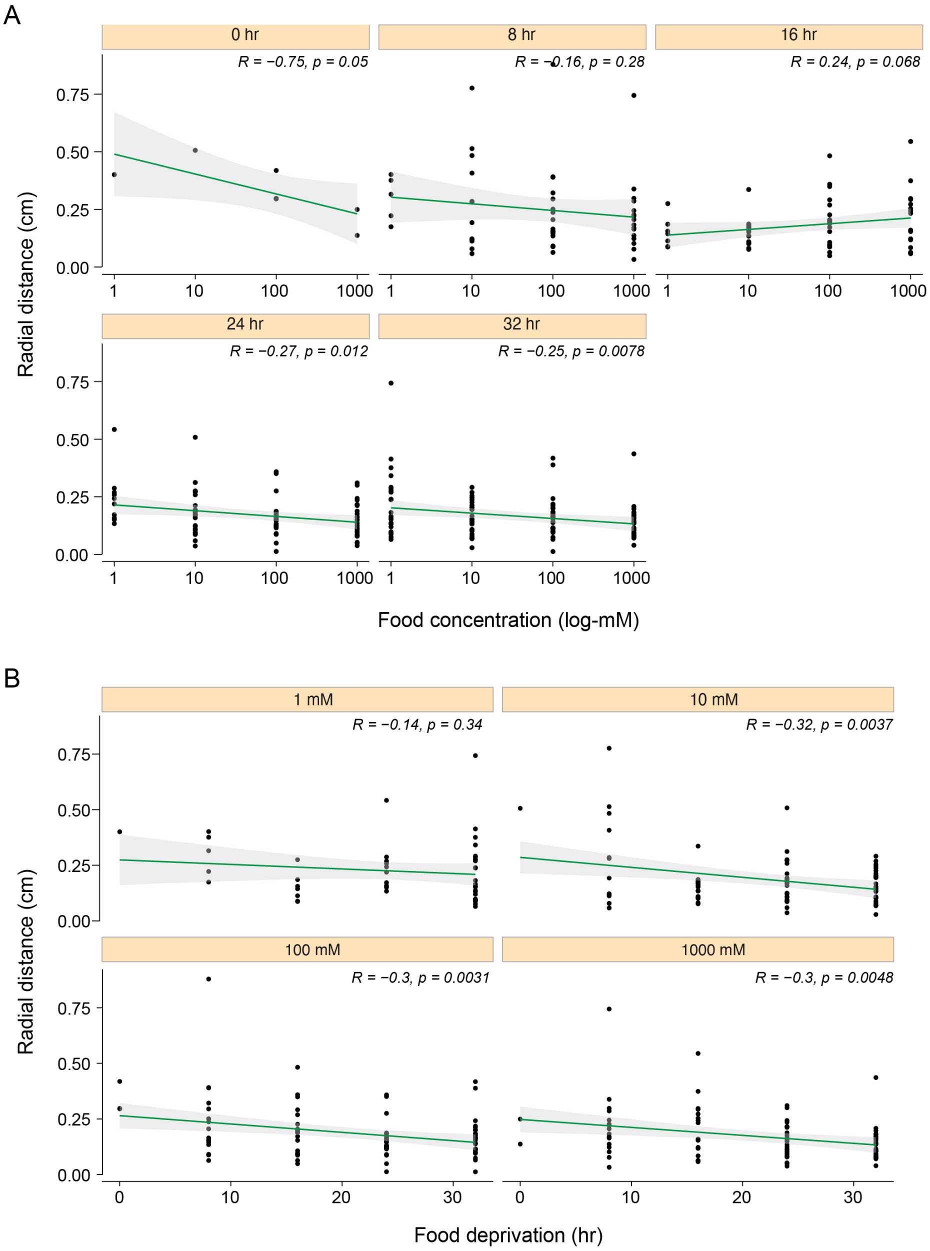


**Figure S3. Metabolic state and taste/nutrient value influence the radial distance of flies to the food source**

**(A)** There is an inverse linear correlation between the radial distance and food concentration. This correlation is not present in flies fed ad libitum or food-deprived for 8 to 16 hours. In contrast, all flies that have been deprived of food for 24 to 32 hours decrease their radial distance from the food source as food concentration rises.

**(B)** There is an inverse linear correlation between the radial distance and food deprivation. This linear relationship is not present at 1mM sucrose solution but is strongly significant at other concentrations of sucrose tested (10mM, 100mM, and 1M).

| Parameter1 | Parameter2 | R | p | Method | n_Obs |
|---|---|---|---|---|---|
| Nutrient value | Radial distance | -0.112263606 | 0.047920204 | Pearson | 311 |
| Nutrient value | Food deprivation | 0.070119785 | 0.217532928 | Pearson | 311 |
| Nutrient value | Meal volume | 0.532581988 | 3.50122E-24 | Pearson | 311 |
| Nutrient value | Food concentration | 0.869897002 | 7.41165E-97 | Pearson | 311 |
| Food concentration | Radial distance | -0.08972159 | 0.114319465 | Pearson | 311 |
| Food concentration | Food deprivation | -0.04865237 | 0.392525995 | Pearson | 311 |
| Food concentration | Meal volume | 0.306195756 | 3.56787E-08 | Pearson | 311 |
| Food concentration | Nutrient value | 0.869897002 | 7.41165E-97 | Pearson | 311 |
| Food deprivation | Radial distance | -0.255821081 | 4.88772E-06 | Pearson | 311 |
| Food deprivation | Food concentration | -0.04865237 | 0.392525995 | Pearson | 311 |
| Food deprivation | Nutrient value | 0.070119785 | 0.217532928 | Pearson | 311 |
| Food deprivation | Meal volume | 0.198174609 | 0.000438586 | Pearson | 311 |
| Meal volume | Radial distance | -0.174710877 | 0.001984971 | Pearson | 311 |
| Meal volume | Food deprivation | 0.198174609 | 0.000438586 | Pearson | 311 |
| Meal volume | Food concentration | 0.306195756 | 3.56787E-08 | Pearson | 311 |
| Meal volume | Nutrient value | 0.532581988 | 3.50122E-24 | Pearson | 311 |
| Radial distance | Food deprivation | -0.255821081 | 4.88772E-06 | Pearson | 311 |
| Radial distance | Meal volume | -0.174710877 | 0.001984971 | Pearson | 311 |
| Radial distance | Nutrient value | -0.112263606 | 0.047920204 | Pearson | 311 |
| Radial distance | Food concentration | -0.08972159 | 0.114319465 | Pearson | 311 |

**Table S1.** The table shows the correlation coefficients between the metabolic and behavioral variables used in the GLM model.

| Model | AICc | Rank |
|---|---|---|
| RD ~ 1 + Food deprivation + Meal Volume | -599.0631208 | 1 |
| RD ~ 1 + Food deprivation + Meal Volume + Food concentration | -598.6174723 | 2 |
| RD ~ 1 + Food deprivation + Meal Volume + Nutrient value | -597.4101306 | 3 |
| RD ~ 1 + Food deprivation + Meal Volume + Food concentration + Nutrient value | -597.3472041 | 4 |
| RD ~ 1 + Food deprivation + Food concentration | -596.0726555 | 5 |
| RD ~ 1 + Food deprivation + Nutrient value | -595.7733567 | 6 |
| RD ~ 1 + Food deprivation + Food concentration + Nutrient value | -594.1664152 | 7 |
| RD ~ 1 + Food deprivation | -593.7182281 | 8 |
| RD ~ 1 + Meal Volume | -581.8153728 | 9 |
| RD ~ 1 + Meal Volume + Food concentration | -580.1968325 | 10 |
| RD ~ 1 + Meal Volume + Nutrient value | -579.9006706 | 11 |
| RD ~ 1 + Meal Volume + Food concentration + Nutrient value | -578.2787232 | 12 |
| RD ~ 1 + Nutrient value | -574.0345331 | 13 |
| RD ~ 1 + Food concentration + Nutrient value | -572.1663853 | 14 |
| RD ~ 1 + Food concentration | -571.943028 | 15 |
| RD ~ 1 | -570.7083634 | 16 |

**Table S2.** The table shows the Gamma GLM models and their ranking according to AICc scores.

| *Figure* | *Abbreviation* | *Full genotype* |
|---|---|---|
| ***1B-1H*** | *CS* | $w^{+}$*/Y; +/+; +/+* |
| ***2A-2H*** | *CS* | $w^{+}$*/* $w^{+}$*; +/+; +/+* |
| ***3A-3K*** | *CS* | $w^{+}$*/Y; +/+; +/+* |
| ***4A-4J*** | *CS* | $w^{+}$*/Y; +/+; +/+* |
| ***5A-5K*** | *+/+* | $w^{1118}$*/Y; +/+; +/+* |
| ***5A-5K*** | *ΔGr5a* | $Gr5a^{LEXA}$*/Y; +/+; +/+* |
| ***5A-5K*** | *ΔGr64f* | $w^{1118}$*/Y; +/+;* $Gr64f^{LEXA}$ |
| ***5A-5K*** | *ΔGr43a* | $w^{1118}$*/Y;* $Gr43a^{GAL4}$*/+; +/+* |
| ***5A-5K*** | *Sugar-blind* | *R1,* $Gr5a^{LexA}$*/Y; ΔGr43a/cyo; Δ61a,Δ64a-f/Tm6b* |
| ***S2*** | *CS* | $w^{+}$*/Y; +/+; +/+* |
| ***S3*** | *CS* | $w^{+}$*/Y; +/+; +/+* |

**Table S3.** The table shows fly genotypes in the main and supplementary figures.
